# PIKFYVE inhibition induces endosome- and lysosome-derived vacuole enlargement via ammonium accumulation

**DOI:** 10.1101/2024.04.19.590364

**Authors:** Junsuke Uwada, Hitomi Nakazawa, Takeshi Kiyoi, Takashi Yazawa, Ikunobu Muramatsu, Takayoshi Masuoka

## Abstract

FYVE-type zinc finger-containing phosphoinositide kinase (PIKFYVE), that is essential for PtdIns(3,5)P_2_ production, is an important regulator of lysosomal homeostasis. PIKFYVE dysfunction leads to cytoplasmic vacuolization; however, the underlying mechanism remains unknown. In this study, we explored the cause of vacuole enlargement upon PIKFYVE inhibition in DU145 prostate cancer cells. Enlargement of vacuoles by PIKFYVE inhibition required glutamine and its metabolism by glutaminases. Addition of ammonia, a metabolite of glutamine, was sufficient to enlarge vacuoles via PIKFYVE inhibition. Moreover, PIKFYVE inhibition led to intracellular ammonium accumulation. Endosome–lysosome permeabilization resulted in ammonium leakage from the cells, indicating ammonium accumulation in the endosomes and lysosomes. Ammonium accumulation and vacuole expansion were suppressed by the lysosomal lumen neutralization. It is therefore assumed that PIKFYVE inhibition interferes with the efflux of NH_4_^+^, which is protonated NH_3_ in the lysosomal lumen, leading to osmotic swelling of vacuoles. Notably, glutamine or ammonium is required for PIKFYVE inhibition-induced suppression of lysosomal function and autophagic flux. In conclusion, this study showed that PIKfyve inhibition disrupts lysosomal homeostasis via ammonium accumulation.

**Summary statement:** Inhibition of the phosphoinositide kinase PIKFYVE results in endosome/lysosome enlargement and impaired lysosomal function. This study showed that the accumulation of glutamine-derived ammonium is the cause of these events.

## Introduction

Lysosomes are acidic intracellular vesicles containing various hydrolytic enzymes, such as cathepsin, that degrade and recycle biomolecules. Lysosomes degrade materials via endocytosis by fusing with endosomes and contribute to macroautophagy/autophagy-induced degradation by fusing with autophagosomes to form autolysosomes (Settembre et al., 2013). Lysosomal dysfunction leads to the accumulation of substrates to be degraded, resulting in various pathologies, such as sphingolipidosis and mucopolysaccharidosis (La Cognata et al., 2020). Lysosomal hydrolases are transported to early endosomes via the trans-Golgi network. Early endosomes are converted into late endosomes, which eventually fuse with lysosomes to maintain their function (Yang and Wang, 2021). Vacuolar H^+^-ATPases (V-ATPases) localize to the membranes of endosomes and lysosomes and maintain their luminal compartments at an acidic pH. Decreased luminal acidity due to V-ATPase inhibition results in decreased hydrolase activity and reduced autophagic flux (Kawai et al., 2007; Yamamoto et al., 1998).

Phosphatidylinositol 3,5-bisphosphate (PtdIns(3,5)P_2_) is a phosphoinositide specifically localized in endosomes and lysosomes (Takatori et al., 2016). FYVE-type zinc finger-containing phosphoinositide kinase (PIKFYVE) is an essential kinase for the production of PtdIns(3,5)P_2_. PIKFYVE localizes to the membranes of endosomes and lysosomes as a complex with 5-phosphatase FIG4 and scaffolding protein VAC14 and phosphorylates PtdIns3P (Hasegawa et al., 2017). Deletion of PIKFYVE results in the loss of intracellular PtdIns(3,5)P_2_ (Zolov et al., 2012). PIKFYVE plays an important role in lysosomal homeostasis by controlling the reformation of lysosomes (Bissig et al., 2017; Yordanov et al., 2019). PIKFYVE is also required for normal autophagic flux, as it is involved in the formation of autolysosomes via the fusion of lysosomes and autophagosomes (Sano et al., 2016; Sharma et al., 2019).

*PIKFYVE*-deficient mice die during embryogenesis (Ikonomov et al., 2011; Takasuga et al., 2013). Studies on PIKFYVE low-expressing and tissue-specific knockout mice have shown that PIKFYVE is essential for the normal development and function of various tissues (Takasuga et al., 2013; Zolov et al., 2012). *PIKFYVE*-deficient tissues showed abnormally swollen LAMP1-positive vacuoles, suggesting that they are endosomes or lysosomes. Genetic abnormalities in *Fig4* or *Vac14* also reduce PtdIns(3,5)P_2_ production, leading to neurodegeneration and enlarged cytoplasmic vacuoles (Chow et al., 2007; Zhang et al., 2007). An in vitro study has shown that deficiency in PIKFYVE kinase-mediated PtdIns(3,5)P_2_ production leads to characteristic vacuolation (Ikonomov et al., 2002). The formation of these vacuoles may be related to dysregulated endosome–lysosome function; however, the underlying mechanism remains unclear. Previous studies have indicated that vacuoles are formed because of an imbalance favoring lysosomal fusion over fission processes (Choy et al., 2018; Sharma et al., 2019). However, whether this abnormality in the lysosomal fusion–fission process was sufficient to explain the vacuole enlargement induced by PIKFYVE inhibition remains unknown.

In this study, we demonstrated that vacuole enlargement induced by PIKFYVE inhibition requires ammonium produced by glutamine metabolism. PIKFYVE inhibition leads to ammonium accumulation in vacuoles. Here, impairment of lysosomal function and autophagic flux due to PIKFYVE inhibition was enhanced by glutamine and ammonium. Our results revealed that vacuoles were produced by PIKFYVE inhibition due to the influx of water associated with the accumulation of ammonium ions in the endosomes and lysosomes. This study provides insights into the mechanism of vacuole expansion during PIKFYVE inhibition and suggests an association between PIKFYVE and glutamine–ammonia metabolism.

## Results

### PIKFYVE inhibition causes endosome and lysosome enlargement

To investigate the effect of PIKFYVE inhibition on cytosolic vacuole formation, DU145 prostate cancer cells were treated with three PIKFYVE inhibitors: YM201636 (YM), apilimod, and SB202190 (SB). Although SB is often used as a p38 mitogen-activated protein kinase (MAPK) inhibitor, it also forms vacuoles via a mechanism other than p38 MAPK inhibition, which has recently been reported to be due to its inhibitory effect on PIKFYVE (Menon et al., 2015; Menon et al., 2011; Wible et al., 2024). Here, treatment with the PIKFYVE inhibitors resulted in the formation of numerous enlarged vacuoles in the cytosol (Fig. 1A,B). These vacuoles were observed as spheres that did not stain well with crystal violet, consistent with a previously report (Menon et al., 2011). PIKFYVE knockdown also induced vacuole formation (Fig. S1A,B). Vacuoles were observed 1 h after YM treatment and gradually expanded over time (Fig. 1C; representative images in Fig. S1C). Early endosome marker EEA was partially localized to several relatively small vesicles formed by YM treatment (Fig. 1D). In contrast, the localization of lysosomal-associated membrane protein 1 (LAMP1) and RAB7A, markers of late endosomes and lysosomes, was consistent with many enlarged vacuoles. Therefore, most vacuoles formed after PIKFYVE inhibition were derived from the late endosomes and lysosomes. Concanamycin B (ConB) suppresses endosome and lysosome acidification by inhibiting V-ATPase (Woo et al., 1992). Chloroquine (CQ), a weak base, also neutralizes the pH by accumulating in lysosomes (Poole and Ohkuma, 1981). Here, treatment with ConB or CQ abolished the YM-induced vacuole enlargement (Fig. 1E). This result suggests that the necessity of an acidic environment in the endosome– lysosome for vacuole formation after PIKFYVE inhibition.

**Fig. 1.**
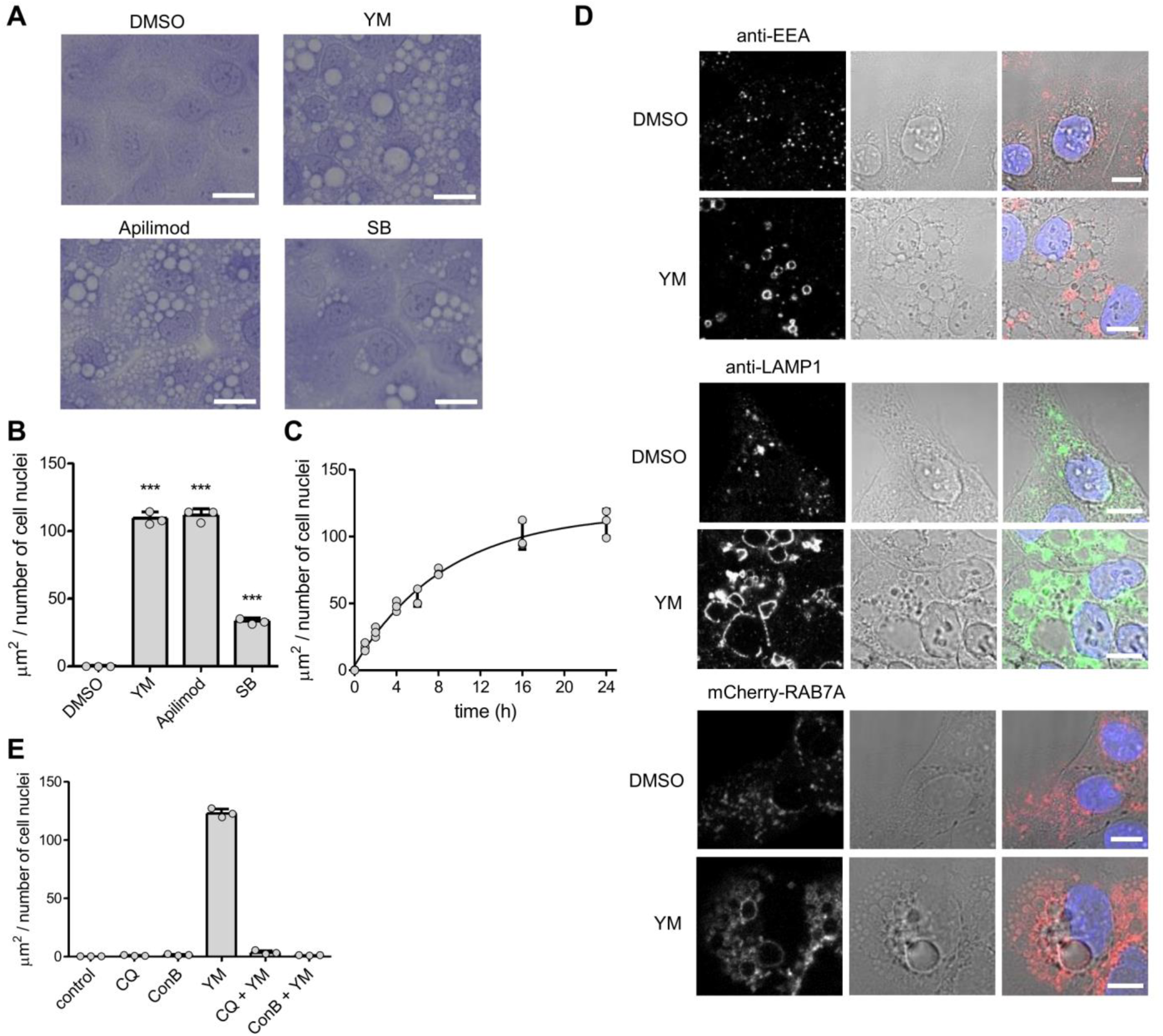
Inhibition of PIKFYVE causes enlarged endosomes and lysosomes. (A and B) DU145 cells were treated with 1 μM YM201636 (YM), 0.03 μM apilimod, 20 μM SB202190 (SB) or dimethyl sulfoxide (DMSO; final: 0.5%) for 24 h. (A) Representative images of crystal violet-stained cells. Bar = 20 μm. (B) Area of vacuoles per nuclei. ***p < 0.001 (compared to DMSO; n = 3). (C) Time-course of vacuole enlargement after 1 μM YM treatment (n = 3). (D) DU145 cells were treated with 1 μM YM or DMSO (final: 0.5%) for 24 h. Immunofluorescence images for EEA or LAMP1, or mCherry-RAB7A (left panels) and bright-field images of cells (middle panels) were merged with DAPI staining (right panels). Bar = 10 μm (E) DU145 cells were co-treated with 1 μM YM and 50 μM chloroquine (CQ) or 0.1 μM concanamycin B (ConB) for 24 h. After crystal violet staining, area of vacuoles was quantified (n = 3).

### Glutamine facilitates PIKFYVE inhibition-induced vacuole enlargement

Vacuolization, which occurs via the expression of a PIKFYVE-binding mutant of *Vac14,* is suppressed by amino acid removal (Schulze et al., 2017). To investigate the relevance of amino acids in vacuolation induced by PIKFYVE inhibition, cells were treated with PIKFYVE inhibitors in an amino acid-free medium in this study. As shown in Fig. 2A, vacuole enlargement by each PIKFYVE inhibitor was suppressed by the amino acid removal. Amino acid depletion leads to the suppression of mechanistic target of rapamycin complex 1 (MTORC1), thus activating autophagy (Takahara et al., 2020). Inhibition of MTORC1 with rapamycin had no effect on vacuole enlargement following PIKFYVE inhibition (Fig. 2B). To determine the specific amino acids critical for vacuole enlargement, we compared the contribution of each amino acid by adding non-essential amino acids (NEAAs) without glutamine, essential amino acids (EAAs), and glutamine to the amino acid-free medium. The presence of glutamine was the most important factor in vacuole enlargement following YM treatment (Fig. 2C; representative images in Fig. S2A). The presence of each amino acid slightly improved cell viability, whereas no critical cell damage was observed, even in the amino acid-free medium culture for 24 h (Fig. S2B). Glutamine is mainly catabolized by glutaminase (GLS) in the mitochondria. Mammals contain two GLS genes: the ubiquitously distributed GLS1 and liver-type GLS2. To examine the involvement of GLS in vacuole enlargement, cells were treated with bis-2-(5-phenylacetamido-1,3,4-thiadiazol-2-yl) ethyl sulfide (BPTES) as a GLS1 inhibitor and glutaminase inhibitor 968 (GI968) as a GLS2 selective inhibitor, as previously described (Lukey et al., 2019). Both BPTES and GI968 inhibited vacuole enlargement, and further inhibition was observed via co-treatment with both inhibitors (Fig. 2D; representative images in Fig. S2C).

**Fig. 2.**
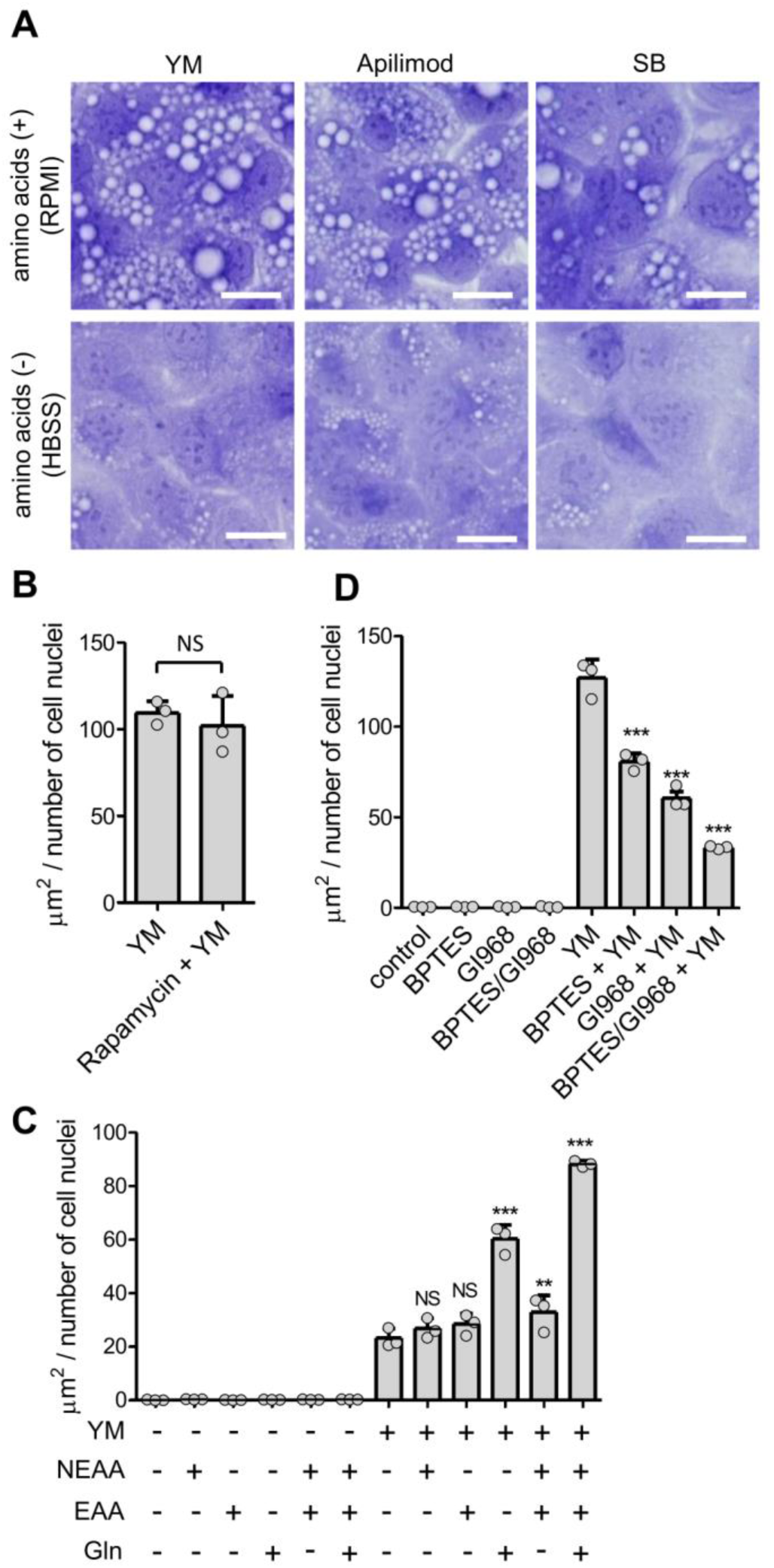
Glutamine facilitates PIKFYVE inhibition-induced vacuole enlargement. (A) DU145 cells were treated with 1 μM YM, 0.03 μM apilimod, and 20 μM SB for 24 h in RPMI-1640 (RPMI) or amino acid-free HBSS medium. Representative images of crystal violet-stained cells. Bar = 20 μm. (B) DU145 cells were co-treated with 1 μM YM and 0.1 μM rapamycin for 24 h in RPMI. After crystal violet staining, area of vacuoles was quantified (n = 3). NS, not significantly different via unpaired two-tail Student’s *t*-test. (C) DU145 cells were treated with 1 μM YM for 24 h in HBSS medium with NEAAs, EAAs or glutamine (Gln). After crystal violet staining, area of vacuoles was quantified. NS, not significantly different. **p < 0.01 and ***p < 0.001 (compared to YM without amino acids; n = 3). (D) DU145 cells were co-treated with 1 μM YM and 10 μM BPTES or 20 μM GI968 for 24 h in RPMI. After crystal violet staining, area of vacuoles was quantified (n = 3). ***p < 0.001 (compared to YM).

Our findings suggest that PIKFYVE inhibition-induced enlargement of vacuoles requires glutamine and its metabolism via GLS.

### Ammonia is necessary for PIKFYVE inhibition-induced vacuole enlargement

Glutamine is catabolized into glutamate and ammonia by GLS in mitochondria (Altman et al., 2016). Glutamate is then converted to α-ketoglutarate (α-KG), which enters the tricarboxylic acid (TCA) cycle. Glutamate is also used in the synthesis of glutathione, which contributes to scavenging of reactive oxygen species. To clarify which of these glutamine metabolites are involved in PIKFYVE inhibition-induced vacuole enlargement, metabolite substitutes were added to the glutamine-depleted medium. As shown in Fig. 3A and B, the addition of ammonium chloride (NH_4_Cl) instead of glutamine was sufficient for vacuole enlargement. Vacuole expansion plateaued at 1 mM NH_4_Cl (Fig. S3A). Similar to CQ, high concentrations of NH_4_Cl are often used as lysosome-neutralizing agents. However, YM-induced vacuole enlargement was not prevented by treatment with 20 mM NH_4_Cl in the RPMI-1640 (RPMI) medium (Fig. S3B). Ammonia scavengers 4-phenylbutyric acid (4-PBA) and methyl pyruvate (MP) suppressed vacuole enlargement in the glutamine-containing culture medium (Fig. 3C; representative images in Fig. S3C).

**Fig. 3.**
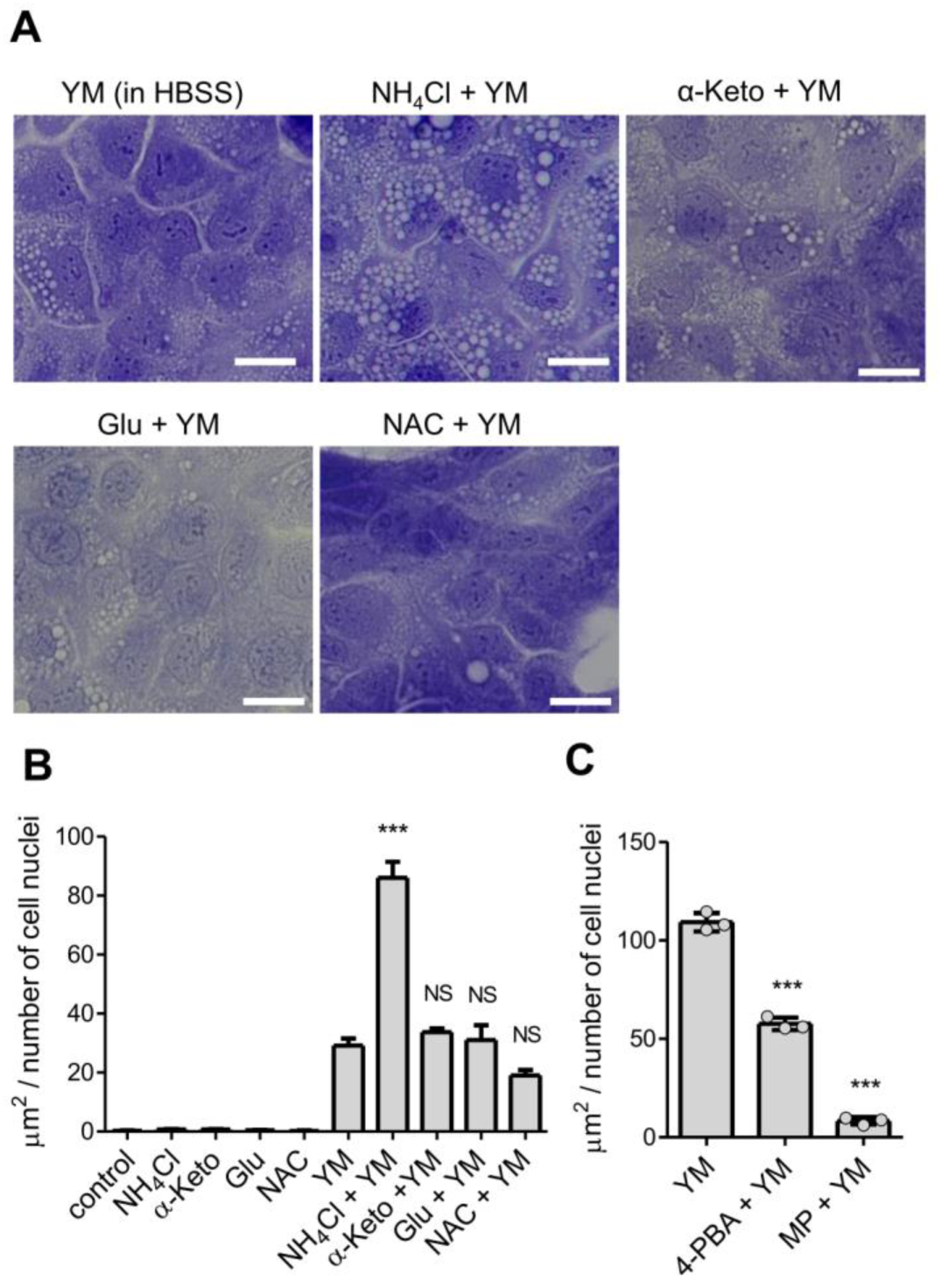
Ammonia is necessary for PIKFYVE inhibition-induced vacuole enlargement. (A and B) DU145 cells were treated with 1 μM YM in HBSS supplemented with 2 mM ammonium chloride (NH_4_Cl), 2 mM dimethyl-2-ketoglutarate (cell-permeable α-ketoglutarate [α-Keto]), 2 mM dimethyl glutamate (cell-permeable glutamate [Glu]) and 10 mM N-acetyl-L-cysteine (NAC) for 24 h. (A) Representative images of crystal violet-stained cells. Bar = 20 μm. (B) Area of vacuoles per nuclei. NS, not significantly different. ***p < 0.001 (compared to YM; n = 3). (C) DU145 cells were co-treated with 1 μM YM and 5 mM 4-PBA or 20 mM methyl pyruvate (MP) for 24 h in RPMI. After crystal violet staining, area of vacuoles was quantified (n = 3). ***p < 0.001 (compared to YM).

Therefore, ammonia derived from glutamine is necessary for vacuole enlargement induced by PIKFYVE inhibition.

### PIKFYVE inhibition leads to ammonium ion accumulation in endosomes and lysosomes

NH_3_ can move across lipid membranes via passive diffusion, whereas NH_4_^+^ requires a transporter to pass through lipid membranes. The mammalian plasma membrane contains the Rhesus (Rh) protein family, which regulates the efflux of intracellularly accumulated NH_4_^+^ (Williamson et al., 2024). The pKa of NH_3_/NH_4_^+^ is approximately 9.25. At neutral pH in the cytosol, NH_3_ is partially present and can therefore, pass through lipid membranes and enter endosomes and lysosomes. However, in endosomes and lysosomes at pH 4.5–6.5, almost all NH_3_ is protonated to NH_4_^+^ and thus cannot leave the vesicle by passive diffusion. Since the enlargement of vacuoles by PIKFYVE inhibition requires ammonia, it was predicted that the accumulation of NH_4_^+^ within the endosomes and lysosomes caused their enlargement. Cellular NH_3_/NH_4_^+^ was evaluated using a modified version of Berthelot reaction (Spinelli et al., 2017). Treatment with the PIKFYVE inhibitors led to the accumulation of cellular NH_3_/NH_4_^+^ (Fig. 4A). PIKFYVE knockdown also induced cellular NH_3_/NH_4_^+^ accumulation (Fig. S4A). NH_3_/NH_4_^+^ increased over time during YM treatment (Fig. 4B). The accumulation of NH_3_/NH_4_^+^ was suppressed by CQ or ConB treatment (Fig. 4C). This result suggests that the acidic environment within endosomes and lysosomes induces the conversion of NH_3_ into NH_4_^+^, resulting in the accumulation of NH_4_^+^ and enlargement of these vesicles. The accumulation of NH_4_^+^ is expected to cause water influx owing to the increased osmotic pressure. Vacuole enlargement by YM was inhibited by exposing the cells to hyperosmotic pressure through the addition of sucrose to the medium (Fig. 4D; representative images in Fig. S4). This indicates that the inhibition of PIKFYVE causes NH_4_^+^ accumulation in endosomes and lysosomes, which leads to water influx and vesicle expansion due to the increase in osmotic pressure.

**Fig. 4.**
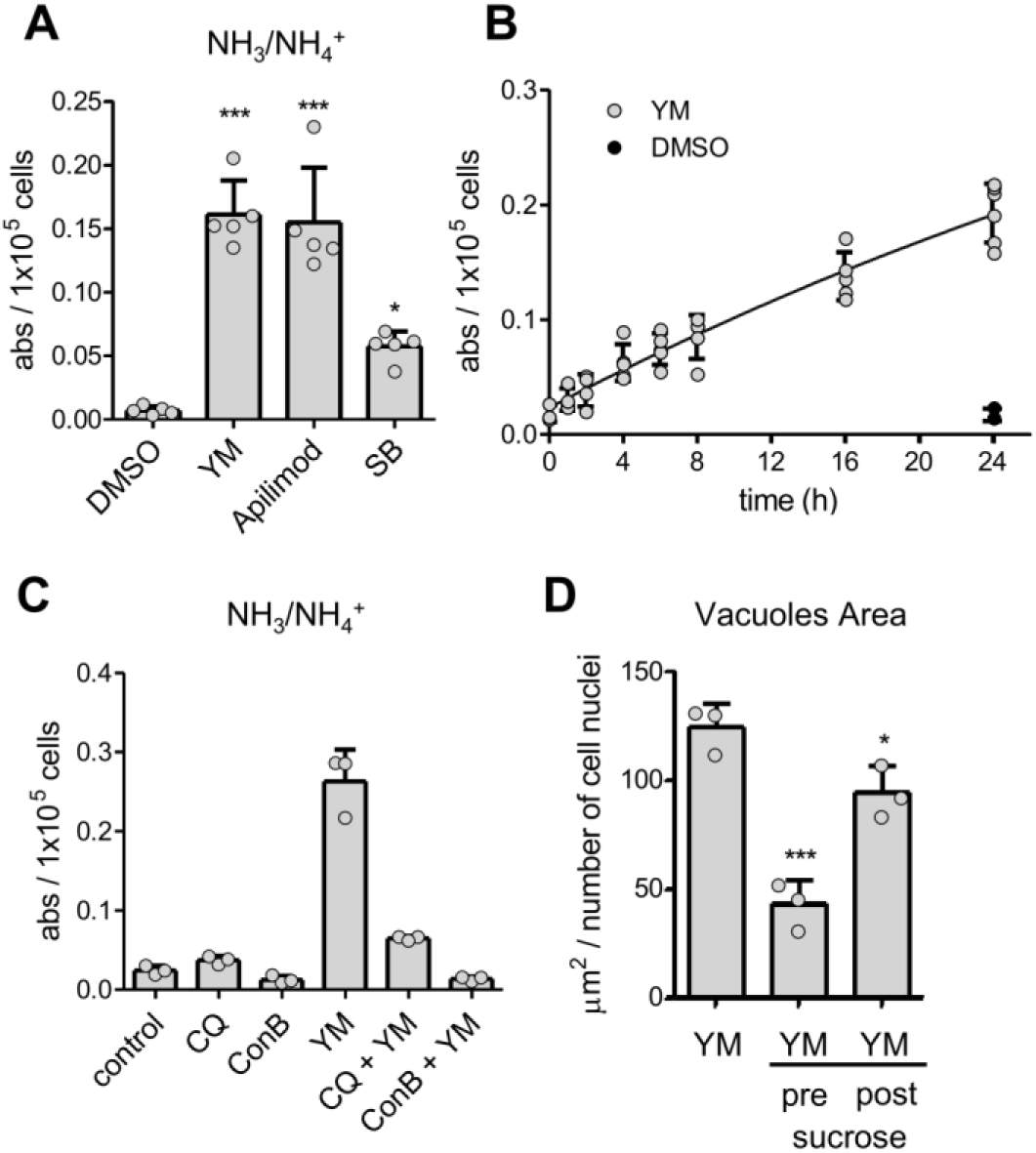
PIKFYVE inhibition leads to ammonium ion accumulation. (A) DU145 cells were treated with 1 μM YM, 0.03 μM apilimod, 20 μM SB or DMSO (final: 0.5%) for 24 h. Each cellular NH_3_/NH_4_^+^ was evaluated using the Berthelot reaction (n = 5). *p < 0.05, ***p < 0.001 (compared to DMSO). (B) Time-course of cellular NH_3_/NH ^+^ accumulation after 1 μM YM or DMSO (final: 0.5%) treatment (n = 3–6). (C) DU145 cells were co-treated with 1 μM YM and 50 μM CQ or 0.1 μM ConB for 24 h. Each cellular NH_3_/NH_4_^+^ was evaluated using the Berthelot reaction (n = 3). (D) DU145 cells were challenged with osmotic stress by the addition of 200 mM sucrose simultaneously with 1 μM YM (pre) or after 24 h YM treatment for 6 h (post). After crystal violet staining, area of vacuoles was quantified (n = 3). *p < 0.05 and ***p < 0.001 (compared to YM).

To confirm that ammonium accumulation occurs in enlarged endosomes and lysosomes, YM-treated cells were exposed to a solution containing graded concentrations of the non-ionic detergent digitonin and the amount of leaked NH_3_/NH_4_^+^ was measured along with the activity of leaked cytosolic lactate dehydrogenase (LDH) and lysosomal cathepsin (Fig. 5A) (Jaattela and Nylandsted, 2015). LDH leakage due to plasma membrane permeabilization was significantly observed at 10 μg/mL of digitonin, reaching approximately 80% of leakage at the final concentration (200 μg/mL) by 20 μg/mL (Fig. 5B). In contrast, cathepsin leakage was not seen until 30 μg/mL and showed a gradual increase from 40 μg/mL. Ammonia leaked gradually from 30 μg/mL digitonin. These results suggest that ammonium accumulates within intracellular organelles, such as endosomes or lysosomes, rather than in the cytoplasm. Next, we examined whether damage to lysosomal membranes affected ammonium accumulation. Injury to the lysosomal membrane with L-leucyl-L-leucine methyl ester (LLOMe) in YM-treated cells caused vacuoles to stain with crystal violet (Fig. 5C). Ammonium accumulation was markedly reduced in these cells, whereas LDH leakage was negligible under the same treatment (Fig. 5D,E). This indicates that LLOMe treatment did not cause critical damage to the cells or leakage of ammonium from the cytoplasm. Ammonium released from the endosomes and lysosomes by LLOMe treatment was possibly reduced in the cells via efflux from the cells or consumption via amino acid metabolism.

**Fig. 5.**
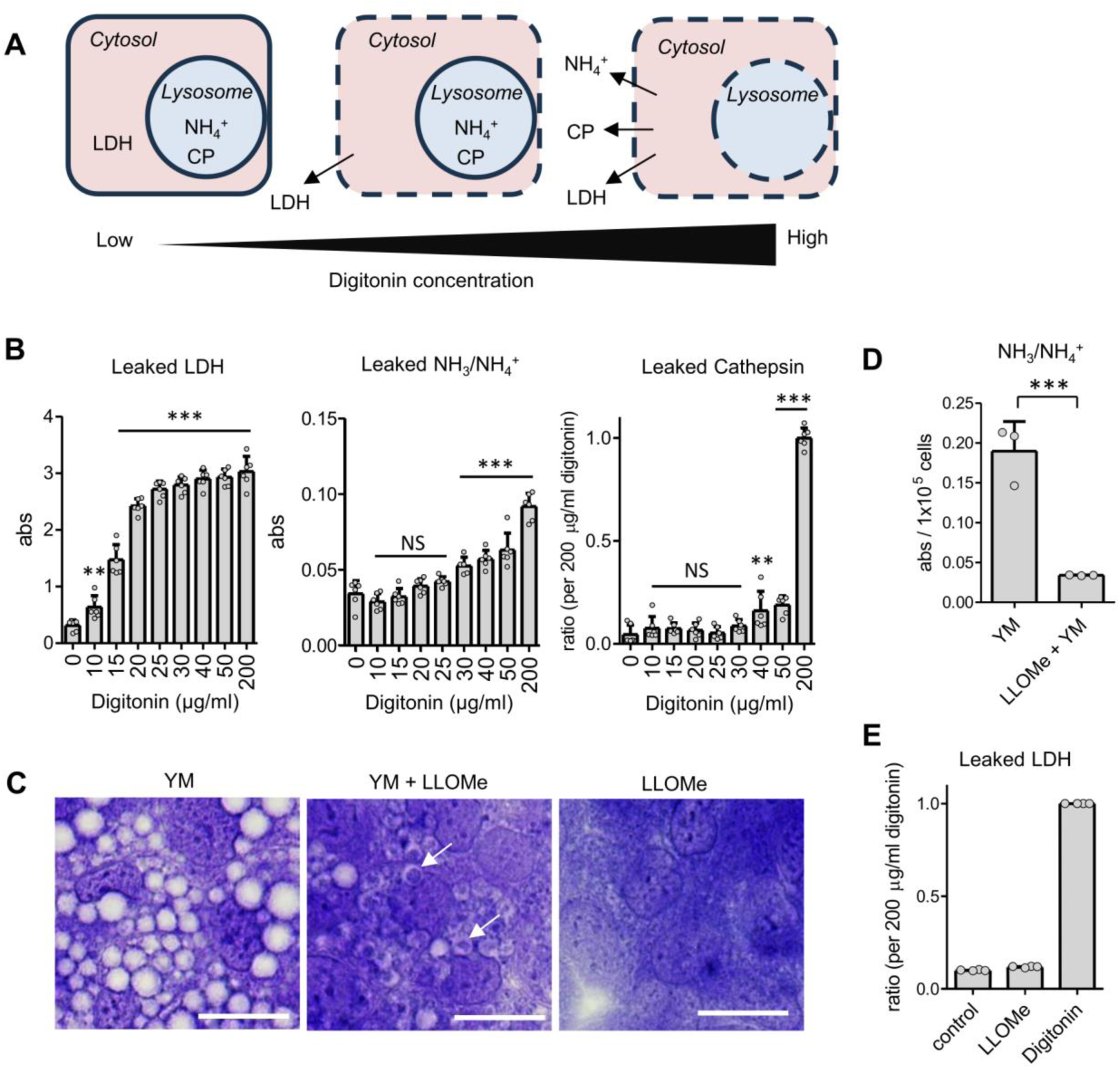
Ammonium accumulation occurs in endosomes and lysosomes during PIKFYVE inhibition. (A) Schematic diagram of the lysosomal permeabilization experiment. Dashed lines represent the lipid membranes with holes in response to increasing digitonin concentrations. CP = cathepsin. (B) Lysosomal permeabilization experiment of DU145 cells. After treating with 1 μM YM for 24 h, DU145 cells were treated with the indicated concentration of digitonin buffer. Leaked NH_3_/NH_4_^+^, LDH, and cathepsin from cells were analyzed (n = 5–6). *p < 0.05, **p < 0.01 and ***p < 0.001 (compared to 0 μg/mL digitonin). (C–E) After 24 h of treatment with or without 1 μM YM, 3 mM LLOMe was applied to the cells. (C) Representative images of crystal violet-stained cells. White arrows indicate the vacuoles that became stained with crystal violet after LLOMe treatment. Bar = 20 μm. (D) Reduction of cellular NH_3_/NH_4_^+^ accumulation by LLOMe treatment for 3 h (n = 3). ***p < 0.001 (unpaired two-tail Student’s *t*-test). (E) Leaked cytosolic LDH after sequential treatment with 3 mM LLOMe for 3 h and 200 μg/mL digitonin for 15 min (n = 4).

### Glutamine and ammonium contribute to the decrease in lysosomal function and autophagic flux via PIKFYVE inhibition

Cathepsins are the most abundant hydrolases, and their dysfunction is associated with various lysosomal storage diseases, such as neuronal ceroid lipofuscinosis 10 (Drobny et al., 2022). Cathepsins are sorted into endosomes as inactive procathepsins, which are eventually processed into active mature forms consisting of light and heavy chains. Inhibition of PIKFYVE suppresses cathepsin D maturation (Sano et al., 2016). Hence, we examined whether glutamine and its metabolite, ammonium, were involved in the maturation of cathepsin D upon PIKFYVE inhibition. As shown in Fig. 6A, suppression of cathepsin D maturation by YM was facilitated by glutamine or ammonium in the medium.

**Fig. 6.**
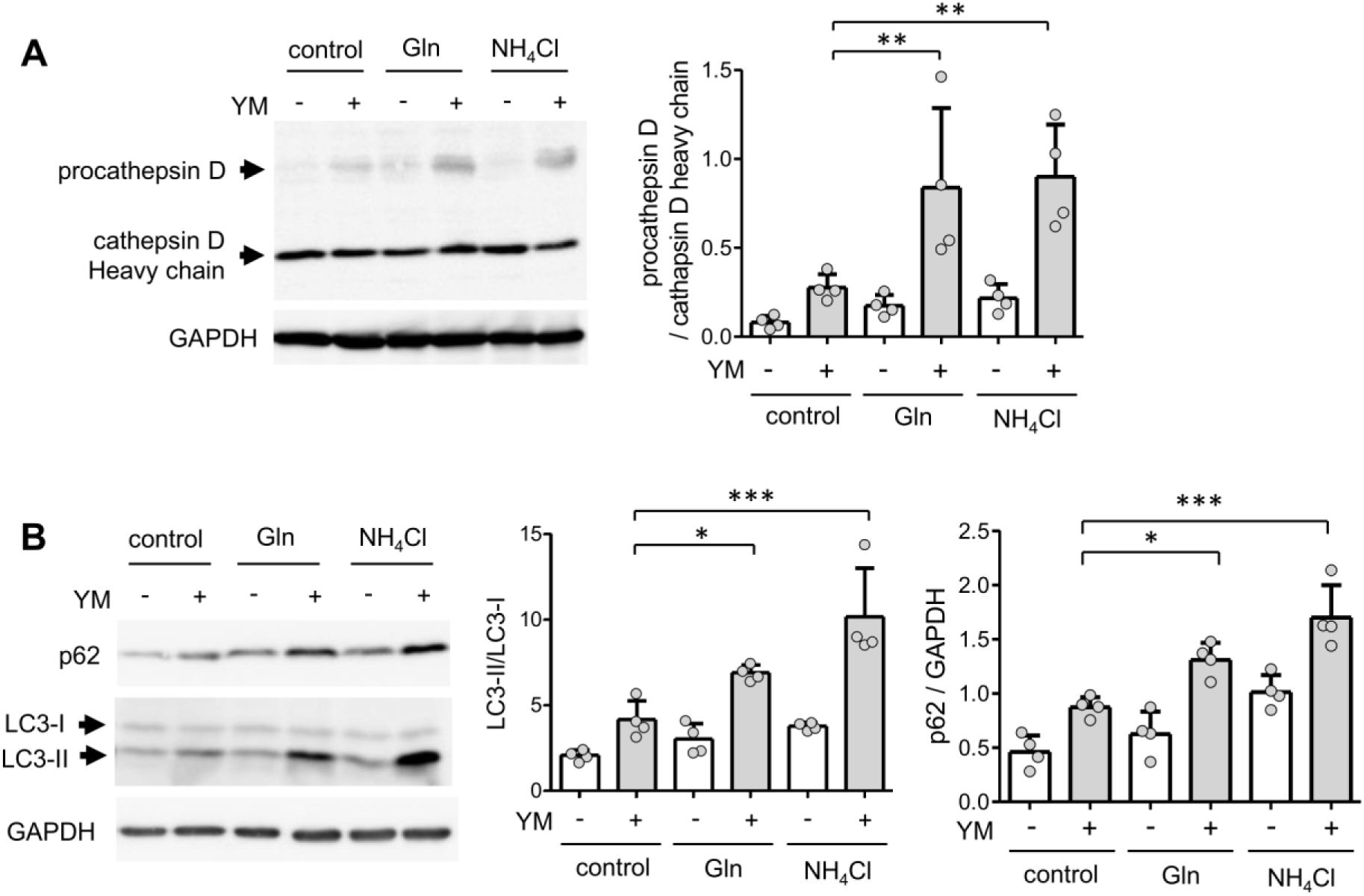
Glutamine and ammonia contribute to the decrease in lysosomal function and autophagic flux via PIKFYVE inhibition. (A) DU145 cells were treated with 1 μM YM for 24 h in HBSS medium containing 2 mM Gln or 2 mM NH_4_Cl. Cell lysates were analyzed via immunoblotting using cathepsin D and GAPDH antibodies. As indicated by the arrows, procathepsin D was observed at high molecular weights, whereas the heavy chain of mature cathepsin D was observed at low molecular weights. The reduced level of mature cathepsin D was quantified as the ratio of procathepsin D to cathepsin D heavy chain (right side). **p < 0.01 (n = 6). (B) HT29 cells were treated with 0.3 μM YM for 24 h in HBSS medium containing 2 mM Gln or 2 mM NH_4_Cl. Cell lysates were analyzed via immunoblotting using p62, LC-3 and GAPDH antibodies. The decrease in autophagic flux was estimated by the increase in the standard autophagy marker, LC3-II, relative to LC3-I and the accumulation of autophagy substrate p62 (right side). *p < 0.05, **p < 0.01, and ***p < 0.001 (n = 4).

Lysosomes are responsible for autophagic degradation through the formation of autolysosomes via fusion with autophagosomes. The inhibition of PIKFYVE decreases autophagic flux by preventing autolysosomes formation (Martin et al., 2013). As DU145 cells lack ATG5, which is required for canonical autophagy, HT29 intestinal epithelial cells were used to evaluate the effect of PIKFYVE inhibition on autophagic flux (Peng et al., 2021). Inhibition of PIKFYVE in HT29 cells resulted in glutamine- and ammonium-dependent vacuole enlargement and intracellular accumulation of ammonium, similar to that observed in DU145 cells (Fig. S5A–C). The suppression of cathepsin D maturation by PIKFYVE inhibition was also facilitated by glutamine or ammonium supplementation in HT29 cells (Fig. S5D). YM treatment-induced increase in microtubule-associated protein 1 light chain 3 (LC3)-II levels, indicating autophagosome accumulation, was also enhanced in the presence of glutamine or ammonium in the medium (Fig. 6B). Concurrently, accumulation of sequestosome 1 (SQSTM1, also known as p62), a selective substrate for autophagic degradation, was enhanced. These results indicate that glutamine and its metabolite, ammonia, are important for the suppression of autophagic flux by PIKFYVE inhibition.

### Involvement of TRPML1 and calcium signaling in PIKFYVE inhibition-induced vacuole enlargement

PtdIns(3,5)P_2_ produced by PIKFYVE, activates mucolipin TRP cation channel 1 (MCOLN1, also known as TRPML1), a cation channel on endosomal and lysosomal membranes (Dong et al., 2010). Loss of function of TRPML1 leads to the enlargement of endosomes and lysosomes (Remis et al., 2014; Venugopal et al., 2007). Here, treatment with mucolipin synthetic agonist 1 (ML-SA1), a TRPML1 agonist, suppressed YM-induced vacuole enlargement (Fig. 7A; representative images in Fig. S6A). Moreover, ammonium accumulation induced by YM was reduced by ML-SA1 (Fig. 7B). Therefore, suppression of TRPML1 activity by reducing PtdIns(3,5)P_2_ production may contribute to ammonium accumulation and endosome and lysosome enlargement during PIKFYVE inhibition.

**Fig. 7.**
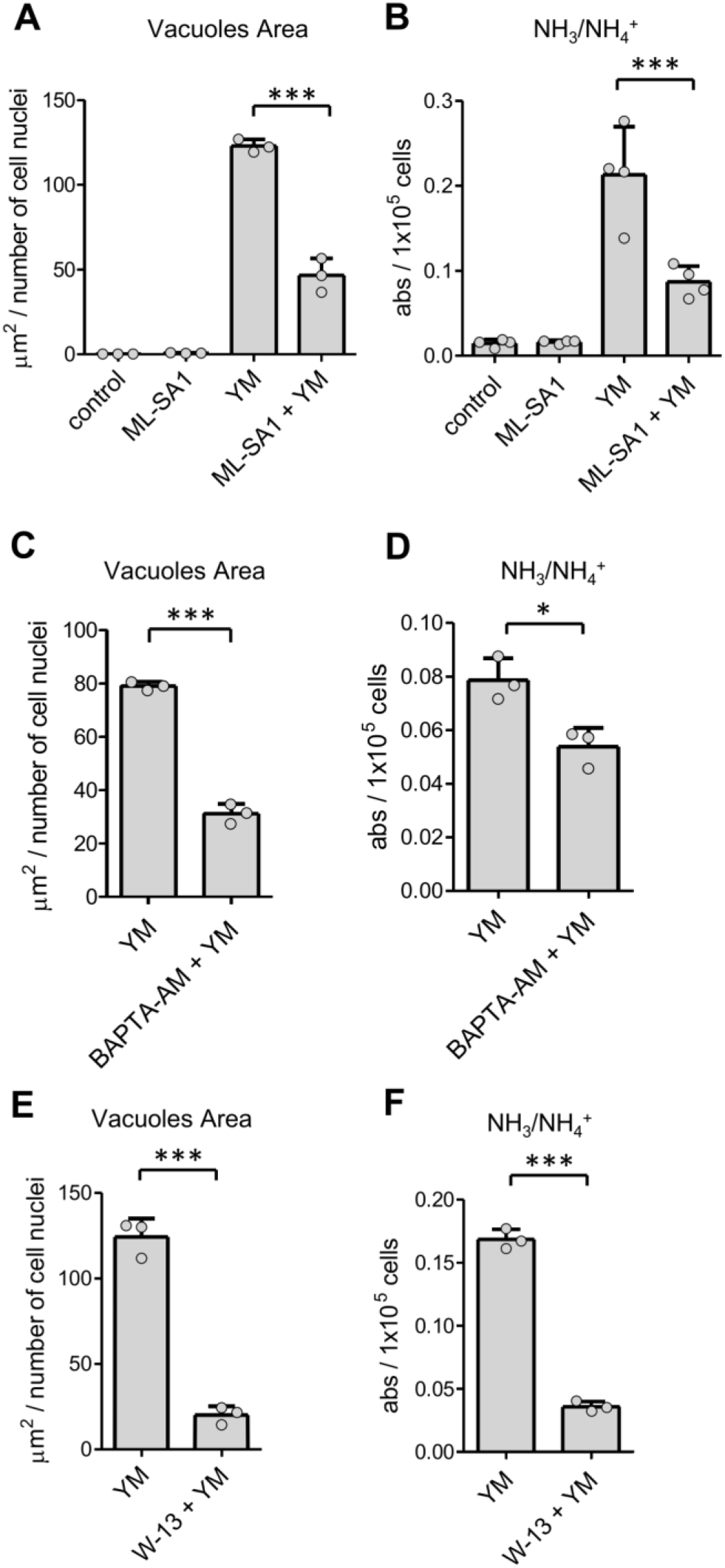
Involvement of TRPML1 and calcium signaling in PIKFYVE inhibition-induced vacuole enlargement. (A and B) DU145 cells were co-treated with 1 μM YM and 20 μM ML-SA1 or DMSO (final: 0.5%) for 24 h. (A) After crystal violet staining, area of vacuoles was quantified. (B) Each cellular NH_3_/NH_4_^+^ was evaluated using the Berthelot reaction. ***p < 0.001 (n = 3). (C and D) DU145 cells were co-treated with 1 μM YM and 20 μM BAPTA-AM or DMSO (final 0.5%) for 6 h. (C) After crystal violet staining, area of vacuoles was quantified. (D) Each cellular NH_3_/NH_4_^+^ was evaluated using the Berthelot reaction. *p < 0.05 and ***p < 0.001 (unpaired two-tail Student’s *t*-test; n = 3). (E and F) DU145 cells were co-treated with 1 μM YM and 20 μM W-13 or DMSO (final: 0.5%) for 24 h. (E) After crystal violet staining, area of vacuoles was quantified. (F) Each cellular NH_3_/NH_4_^+^ was evaluated using the Berthelot reaction. ***p < 0.001 (unpaired two-tail Student’s *t*-test; n = 3).

Finally, the roles of calcium and calmodulin signaling in the changes induced by PIKFYVE inhibition were evaluated. Ca^2+^ and calmodulin are essential for the homotypic and heterotypic fusion of endosomes and lysosomes (Colombo et al., 1997; Pryor et al., 2000). Application of the Ca^2+^ chelator O,O’-Bis(2-aminophenyl)ethyleneglycol-N,N,N,N’-tetraacetic acid, tetraacetoxymethyl ester (BAPTA-AM) or the calmodulin inhibitor W-13 suppressed YM-induced vacuole enlargement and ammonium accumulation (Fig. 7C–F; representative images in Fig. S6B,C). These results suggest that Ca^2+^ and calmodulin contribute not only to vesicle fusion but also to ammonium accumulation, thereby promoting endosome and lysosome enlargement.

## Discussion

In this study, we showed that the enlargement of endosomes and lysosomes by PIKFYVE inhibition was induced by the accumulation of glutamine-derived ammonium. As NH_3_ passes through the lipid membrane via passive diffusion, it penetrates the endosomes and lysosomes. However, in a luminal acidic environment, NH_3_ is protonated to NH_4_^+^ and cannot freely pass through the lipid membrane into the cytosol. Here, neutralization of endosomes and lysosomes with CQ or ConB strongly suppressed ammonium accumulation and vacuole enlargement, suggesting that the protonation of NH_3_ to NH_4_^+^ in an acidic environment is crucial for these events. A high concentration of NH_4_Cl was used as the lysosome neutralizer. However, vacuole enlargement induced by PIKFYVE inhibition was not suppressed by 20 mM NH_4_Cl (Fig. S3B). This result indicated that lysosomal neutralization by NH_4_Cl requires the active efflux of NH_4_^+^ from the lysosomal lumen, which may be blocked by PIKFYVE inhibition. A model of PIKFYVE inhibition-induced lysosomal enlargement and the targets of the drugs used in this study are summarized in a schematic diagram (Fig. S7).

Efflux of accumulated NH_4_^+^ in endosomes and lysosomes requires a specific transporter. Recently, solute carrier family 12 member 9 (SLC12A9) was reported as an ammonium-chloride co-transporter that effluxes lysosomal NH_4_^+^ into the cytoplasm (Levin-Konigsberg et al., 2023). *SLC12A9* knockout causes osmotic swelling of lysosomes via ammonium accumulation. Recently, it was reported that vacuole formation in *FIG4* knockout cells is attenuated by the overexpression of SLC12A9 (Accogli et al., 2024). This suggests that inhibition of SLC12A9-mediated NH_4_^+^/Cl^-^ efflux is involved in vacuole formation during PIKFYVE inhibition. An attractive hypothesis is that under normal conditions, ammonia exchange between mitochondria and lysosomes may control the lysosomal Cl^-^ concentration, thereby coupling glutamine metabolism and lysosomal function. As SLC12A9 requires luminal Cl^-^ for ammonium efflux, PIKFYVE may indirectly regulate SLC12A9 by controlling vesicular Cl^-^ concentrations. Chloride channel (ClC)-7 is a chloride/proton antiporter in endosomes and lysosomes. PtdIns(3,5)P_2_ generated by PIKFYVE inhibits ClC-7. Conversely, inhibition of PIKFYVE promotes ClC-7 activity (Leray et al., 2022). Inhibition of PIKFYVE leads to enlargement of endosomes and lysosomes along with luminal hyperacidification and that these changes are exerted by ClC-7 activation (Cao et al., 2023; Leray et al., 2022). In this case, as the luminal concentration of Cl^-^ should be increased by the activation of ClC-7, the ammonium efflux function of SLC12A9 should not be suppressed by the loss of Cl^-^. Thus, it is plausible that PIKFYVE inhibition may affect SLC12A9 function by mechanisms other than regulating luminal Cl^-^. In contrast, knockout mice of ClC-7 show vacuolated endosomes and lysosomes (Wartosch et al., 2009). Therefore, it remains possible that a decrease in luminal Cl^-^ concentration may affect vacuole enlargement and ammonium accumulation. Future studies should assess the effects of PIKFYVE inhibition on Cl^-^ concentrations in endosomes and lysosomes.

Since accumulation of ammonium in endosomes and lysosomes would be expected to neutralize the luminal pH, the results of previous reports that PIKFYVE inhibition promotes endosomal and lysosomal acidification appear to be contradictory (Cao et al., 2023; Leray et al., 2022). However, once ammonium accumulation reduces the acidity of the vesicles, no further ammonium accumulation or vacuole enlargement is expected to occur, as in the case of V-ATPase inhibition. Our results showed that PIKFYVE inhibition continuously accumulated ammonium over 24 h. Thus, ClC-7-mediated acidification of endosomes and lysosomes may outweigh neutralization by ammonium, leading to persistent ammonium accumulation.

PtdIns(3,5)P_2_ directly activates TRPML1. Vacuole formation in *Vac14* knockout cells is suppressed by TRPML1 overexpression (Dong et al., 2010). Additionally, *Trpml1* knockout leads to enlarged vacuoles (Remis et al., 2014; Venugopal et al., 2007). In this study, we showed that PIKFYVE inhibition-induced vacuole enlargement and ammonium accumulation were reduced by the TRPML1 agonist, ML-SA1. TRPML1 localizes to the endosome–lysosome and contributes to lysosomal homeostasis by controlling vesicular trafficking and lysosomal reformation (Venkatachalam et al., 2015). Disruption of TRPML1 causes lysosomal acidification (Miedel et al., 2008; Soyombo et al., 2006). TRPML1 can be activated by sensing the reduced lysosomal Cl^-^ concentration (Lee and Hong, 2023). Therefore, TRPML1 and ClC-7 may collaborate to maintain lysosomal function, and inhibition of PIKFYVE may impair the lysosomal system by affecting both channels.

Previously, it was proposed that vacuole expansion due to PIKFYVE inhibition may be due to a shift in the fusion/fission balance of endosomal and lysosomal vesicles toward fusion dominance (Choy et al., 2018). Heterotypic and homotypic fusion of endosomes and lysosomes requires Ca^2+^ (Pryor et al., 2000). The results of this study, in which endosome and lysosome enlargement by PIKFYVE inhibition was suppressed by Ca^2+^ depletion and calmodulin inhibition, are consistent with the previous hypothesis.

However, it has also been suggested that this vacuole enlargement is due to osmotic swelling caused by an increase in luminal solute concentration (Cao et al., 2023). Overexpression and activation of the endosomal–lysosomal Ca^2+^ channel P2X4 leads to endosome and lysosome enlargement through the induction of their fusion; however, this enlargement is saturated 1 h after activation (Cao et al., 2015). After PIKFYVE inhibition, enlarged vacuoles were also observed within 1 h, and enlargement continued for 24 h. Simultaneously, ammonium accumulation increased by approximately 14-fold after 24 h compared to 1 h. Our results also show that Ca^2+^ and calmodulin are required for ammonium accumulation. These results suggest that Ca^2+^ and calmodulin contribute to vesicle fusion and ammonium accumulation, resulting in an increase in the luminal solute concentration and osmotic swelling of endosomes and lysosomes.

Decreased PtdIns(3,5)P_2_ production due to PIKFYVE dysfunction causes various pathological disturbances. Fleck corneal dystrophy (FCD) is an autosomal dominant genetic disease characterized by vacuolar swelling in corneal cells that is caused by *PIKFYVE* (Kawasaki et al., 2012; Li et al., 2005). Patients with Mucolipidosis type IV, caused by genetic defects in *TRPML1*, a receptor for PtdIns(3,5)P_2_, exhibit vacuole accumulation in various organs (Wakabayashi et al., 2011). Here, our results revealed that vacuole enlargements in these diseases may be caused by the accumulation of glutamine-derived ammonium. We also showed that the reduction in lysosomal function and autophagic flux by PIKFYVE inhibition requires glutamine or ammonium. Therefore, interference with glutamine–ammonia metabolism may alleviate the symptoms associated with these genetic abnormalities. Brain vacuolation is a characteristic feature of prion disease. PIKFYVE is degraded through deacylation in prion disease, resulting in spongiform neurodegeneration (Lakkaraju et al., 2021). Our results indicate that vacuolation in prion diseases may also be due to the accumulation of ammonium.

PIKFYVE is a potential clinical target for several diseases, including neurodegenerative diseases, such as amyotrophic lateral sclerosis (ALS) and Alzheimer disease, cancer, and viral infections, such as coronavirus disease 2019 (COVID-19) (Burke et al., 2023). Most of its action mechanisms are explained by its effects on lysosomal function and autophagy. Our study showed that glutamine and ammonium contribute to a reduction in lysosomal function and autophagic flux upon PIKFYVE inhibition, suggesting that glutamine–ammonia metabolism in therapeutic target cells may influence the effects of these drugs.

This study demonstrated that ammonia derived from glutamine metabolism can be sequestered in intracellular compartments by pharmacological inhibition of PIKFYVE. Excessive ammonia is toxic; therefore, the liver exerts the urea cycle to maintain low serum ammonia concentration. In hepatic encephalopathy, ammonia metabolism decreases with the liver function, leading to elevated blood ammonia levels and impaired brain function (Braissant et al., 2013). The effect of ammonia sequestration in endosomes and lysosomes via PIKFYVE inhibition on blood ammonia levels and its impact on the pathology of hyperammonemia warrant future investigation. Increased intracellular ammonia levels promote glutamate synthesis, resulting in the depletion of α-KG, a substrate of the TCA cycle. The resulting decrease in ATP synthesis contributes to the abnormal neurotransmission caused by hyperammonemia (Drews et al., 2020). Future studies should investigate whether the sequestration of ammonia in endosomes and lysosomes decreases the cytosolic ammonia levels. A reduction in cytosolic ammonia may also affect amino acid metabolism, such as glutamine synthesis. Cancer cells depend on glutamine and its metabolites for growth (Jin et al., 2023). Therefore, the effects of PIKFYVE inhibitors on glutamate–ammonia metabolism may be associated with their efficacy as anticancer drugs.

In conclusion, this study revealed that the enlargement of endosome- and lysosome-derived vacuoles by PIKFYVE inhibition was due to ammonium accumulation. Our results provide important insights into the diseases associated with PIKFYVE dysfunction. Further investigation of the relationship between PIKFYVE and ammonium transporters is necessary to determine cause of ammonium accumulation in endosomes and lysosomes.

## Materials and Methods

### Materials

All compounds used in this study were from commercial sources. YM201636 (13576), apilimod (19094), SB202190 (21201), BPTES (19284), glutaminase inhibitor compound 968 (17199), rapamycin (13346), and LLOMe (16008) were purchased from Cayman Chemical. Ammonium chloride (015-02991), ML-SA1 (131-18531), MEM NEAA (139-15651), MEM EAA (132-15641), and 200 mM L-glutamine solution (073-05391) were obtained from FUJIFILM Wako Pure Chemical Corporation (Wako). Concanamycin B (BLK-1160) was purchased from BioLinks KK; Dimethyl α-ketoglutarate (K0013), dimethyl L-glutamate hydrochloride (D3353), methyl pyruvate (P0580), digitonin D0540), and W-13 hydrochloride (A2416) were purchased from Tokyo Chemical Industry. Chloroquine diphosphate (08660-04), crystal violet (09804-52), N-acetyl-L-cysteine (00512-84), and 4-(2-Aminoethyl)-benzolsulfonylfluorid-hydrochloride (AEBSF; 02068-64) were obtained from Nacalai Tesque. Sodium 4-phenylbutyrate (4-PBA; P2815) was obtained LKT Laboratories. BAPTA-AM (B035), and 4’,6-diamidino-2-phenylindole, dihydrochloride (DAPI; D212) were purchased from Dojindo. benzyloxycarbonyl-L-phenylalanyl-L-arginine 4-methylcoumaryl-7-amide (Z-FR-MCA; 3095-v) was obtained from the Peptide Institute.

### Cell culture and transfection

DU145 and HT29 cells were maintained in RPMI-1640 medium (Nacalai, 30264-56) supplemented with 10% fetal bovine serum and antibiotics (penicillin and streptomycin; Nacalai, 26253-84). The cells were grown in 5% CO_2_ at 37°C. DU145 cells were kindly provided by Prof. Toshiyuki Kamoto (Department of Urology, Faculty of Medicine, University of Miyazaki) (Kamibeppu et al., 2018). HT29 cells were obtained from the American Type Culture Collection (ATCC). These cells were routinely checked for Mycoplasma infection by DAPI staining. Before treatment with PIKFYVE inhibitors (YM, apilimod, and SB), the medium was replaced with the serum-free RPMI-1640 medium. Other compounds (ConB, CQ, rapamycin, BPTES, GI968, 4-PBA, MP, ML-SA1, and W-13) were applied simultaneously with the PIKFYVE inhibitors and incubate for 24 h. The co-treatment time of BAPTA-AM and YM was shortened to avoid cell detachment (6 h in Fig. 7C,D). As shown in Fig. 5C–E, cells treated with YM for 24 h were challenged with 3 μM LLOMe for 3 h to permeabilize the lysosomal membrane. To estimate the roles of amino acids, the cell culture medium was replaced with the Hank’s balanced salt solution (HBSS; Wako, 084-08345) supplemented with 1.5 mM CaCl_2_, 1.3 mM MgCl_2_, 19.6 mM NaHCO_3_, MEM Vitamin Solution (Wako, 130-17141) and antibiotics and mixed with NEAA, EAA or 2 mM glutamine. The addition of NaHCO_3_ allowed the pH of HBSS to remain neutral in a CO_2_ incubator after 24 h of incubation (pH 7.4). As shown in Fig. 3A, 2 mM dimethyl α-ketoglutarate, 2 mM dimethyl L-glutamate hydrochloride, 2 mM NH_4_Cl or 10 mM N-acetyl-L-cysteine (NAC) was added into HBSS supplemented with NEAA and EAA. Transient transfection was performed using the Xfect Transfection Reagent (Takara Bio, 631316) according to the manufacturer’s instructions. For the knockdown experiments, siRNA against PIKFYVE gene (mixture of Silencer Select Pre-designed siRNA s47254 and s532910; Invitrogen) or control siRNA (AllStar Neg. control siRNA, QIAGEN) was introduced into the DU145 cells. Each oligonucleotide was transiently transfected at a final concentration of 10 nM using Lipofectamine RNAiMAX (Invitrogen) according to the manufacturer’s instructions.

### Crystal violet staining

The cells were seeded on coverslips and cultured for 2–3 d. Then, the cells were treated with PIKFYVE inhibitors for 24 h or the indicated time in a serum-free medium and fixed in glyoxal solution with pH adjusted to 4.0 using NaOH (Richter et al., 2018). For PIKFYVE knockdown, DU145 cells were incubated for 7 d after siRNA transfection and cultured in serum-depleted RPMI medium for 24 h before glyoxal fixation. After washing with water, the cells were stained with 0.05% crystal violet solution for 10 min. Coverslips were washed and mounted on glass slides using phosphate-bufferd saline (PBS). Bright-field images were acquired using the BZ-X700 All-in-One inverted fluorescence microscope (Keyence, Osaka, Japan). Area of vacuoles was analyzed with Fiji (Image J) software. Extraction of vacuoles as regions of interest (ROIs) was performed manually after processing using the Trainable Weka Segmentation plug-in (Arganda-Carreras et al., 2017). Total area of the vacuoles was normalized to the number of cell nuclei per image. Three images were captured per coverslip, and the average of the analyzed results for each image was considered as one experiment. Each result was based on at least three independent experiments.

### Quantitative PCR

The knockdown of PIKFYVE was confirmed using quantitative PCR. After siRNA transfection, the DU145 cells were incubated for three days. cDNA was prepared as previously described (Uwada et al., 2020). Real-time PCR was performed using Thunderbird SYBR qPCR Mix (Toyobo, QPS-101) and QuantStudio 12K Flex Real-Time PCR System (Invitrogen), according to the manufacturer’s instructions. Primers used in this study are as follows: 5’-GACCACTGAGGATGAACGCA-3’ and 5’-GCGGTGATCCTGAAACTCCA -3’ for human PIKFYVE, 5’-CCACTCCTCCACCTTTGACG -3’ and 5’-CACCCTGTTGCTGTAGCCAA -3’ for human glyceraldehyde-3-phosphate dehydrogenase (GAPDH).

### Immunofluorescence assay

DU145 cells grown on coverslips were fixed with 4% paraformaldehyde in PBS for 15 min at room temperature and permeabilized with 0.2% TritonX-100 for 5 min. After permeabilization, the cells were blocked with 0.2% bovine serum albumin and incubated overnight at 4°C with the anti-EEA (1:200; Cell Signaling Technology, 3288) or anti-LAMP1 (1:200; BD Pharmingen, 555798) antibody. The cells were subsequently incubated with the anti-mouse IgG DyLight488 (1:250; Jackson ImmunoResearch, 715-485-151) or anti-rabbit IgG Cy3 (1:250; Jackson ImmunoResearch, 711-165-152) antibody. Subcellular localization of RAB7A was visualized via exogenous expression of mCherry-tagged RAB7A. *RAB7A* was cloned from the cDNA synthesized from total human brain RNA (Khan et al., 2015) into the pmCherry-N1 plasmid (Clontech Laboratories, 632523). Cells were stained with DAPI, and coverslips were mounted on glass slides using glycerol-DABCO. Images were captured using the LSM710 confocal microscope (Carl Zeiss, Jena, Germany).

### Berthelot reaction

Cellular accumulation of ammonia/ammonium was evaluated using a modified version of the Berthelot reaction (Spinelli et al., 2017). Briefly, cells were seeded in 35-mm culture dishes. For PIKFYVE knockdown, DU145 cells were incubated for 7 d after siRNA transfection and then cultured in serum-depleted RPMI medium for 24 h. After PIKFYVE inhibition, cells were detached with 250 μL trypsin solution (Nacalai, 32777-44) and collected in 1.5 mL tubes with 1 mL culture medium. Then, 50 μL of cell suspension was aliquoted for cell counting. Remaining 1200 μL of cell suspension was centrifuged, washed with PBS, centrifuged, and the precipitate was suspended in 100 μL of 80% methanol. After keeping at –30°C for 10 min, the cell suspensions were centrifuged and 15 μL of supernatants were applied to a 96-well dish with 75 μL of solution A (100 mM phenol, 50 mg/L sodium nitroprusside). Next, 75 μL of solution B (0.38 M dibasic sodium phosphate, 125 mM NaOH, and 1% sodium hypochlorite) was added to each well and incubated for 40 min at 37°C. Optical absorbance (Abs) of the aliquots was measured at 635 nm using a microplate reader (TECAN, Infinite M Nano, Männedorf, Switzerland). Abs was subtracted from the background value (80% methanol) and normalized to the number of cells.

### Lysosomal membrane permeabilization assay

Accumulation of ammonium in lysosomes was examined based on the difference in lipid membrane permeability to digitonin between the plasma and lysosomal membranes. The protocol was based on a previously reported method (Jaattela and Nylandsted, 2015). Briefly, cells seeded on the 24 well dish were treated with YM for 24 h. Then, cells were washed with cold PBS and treated with 200 μL of digitonin extraction buffer (250 mM sucrose, 10 mM KCl, 1.5 mM MgCl_2_, 1 mM EDTA, 1 mM EGTA, 0.5 mM AEBSF, 20 mM HEPES, pH 7.5) containing the indicated digitonin concentrations. After rocking on ice for 15 min, the supernatants were collected and centrifuged. Ammonium leakage was analyzed using the Berthelot reaction with addition of 75 μL of Solution A and Solution B. LDH leakage was analyzed using the Cytotoxicity LDH Assay Kit-WST (Dojindo, 347-91751), according to the manufacturer’s instructions. Cathepsin activity was evaluated based on the kinetics of Z-FR-MCA hydrolysis. The supernatants were mixed with the same volume of cathepsin reaction buffer (50 mM sodium acetate, 4 mM EDTA, 0.5 mM AEBSF, 8 mM dithiothreitol, and 50 μM Z-FR-MCA [pH 6.0]). Fluorescence transition (excitation: 380 nm; emission: 460 nm) was measured at 45 s intervals for 20 min using a fluorescence microplate reader (Fluoroskan Ascent FL, Thermo Labsystems, Waltham, MA, USA).

### Western blotting

Samples were prepared and transferred to polyvinylidene difluoride (PVDF) membranes (Wako, 033-22453) as previously described (Uwada et al., 2021). The membranes were probed with the appropriate concentrations of primary antibodies against cathepsin D (1:100; Cell Signaling Technology, 2284), GAPDH (1:200; Wako, 016-25523), p62 (1:1000; MBL, M162-3), and LC3B (1:400; MBL, PM036).

Immunoreactive proteins were detected using a horseradish-peroxidase-labeled secondary antibody for mouse IgG (1:10,000; Jackson ImmunoResearch, 115-035-003) or for anti-rabbit IgG (1:10,000; Jackson ImmunoResearch, 111-035-003) with EzWestLumi plus (ATTO, WSE-7120S). The signal intensity was calculated using the Fiji software. Whole images of western blotting are shown in Supplemental Materials (Fig. S8).

### Statistical analyses

Data were analyzed using the Prism software (GraphPad Software, San Diego, CA, USA). Data are expressed as mean ± standard deviation. The original data plots were superimposed on each graph. All data were analyzed using the one-way analysis of variance followed by Dunnett’s *post-hoc* test, unless stated otherwise in the figure legends. Statistical significance was set at p < 0.05.

## Acknowledgments

The authors thank Hiroko Matsui for the excellent secretarial assistance. All imaging was performed at the Research Support Center of Medical Research Institute in Kanazawa Medical University.

## Data and resource availability

All relevant data and resources can be found in the article and supplementary information.

