## Supplemental Figs for "PIKFYVE inhibition induces endosome- and lysosome-derived vacuole enlargement via ammonium accumulation"

### Fig. S1

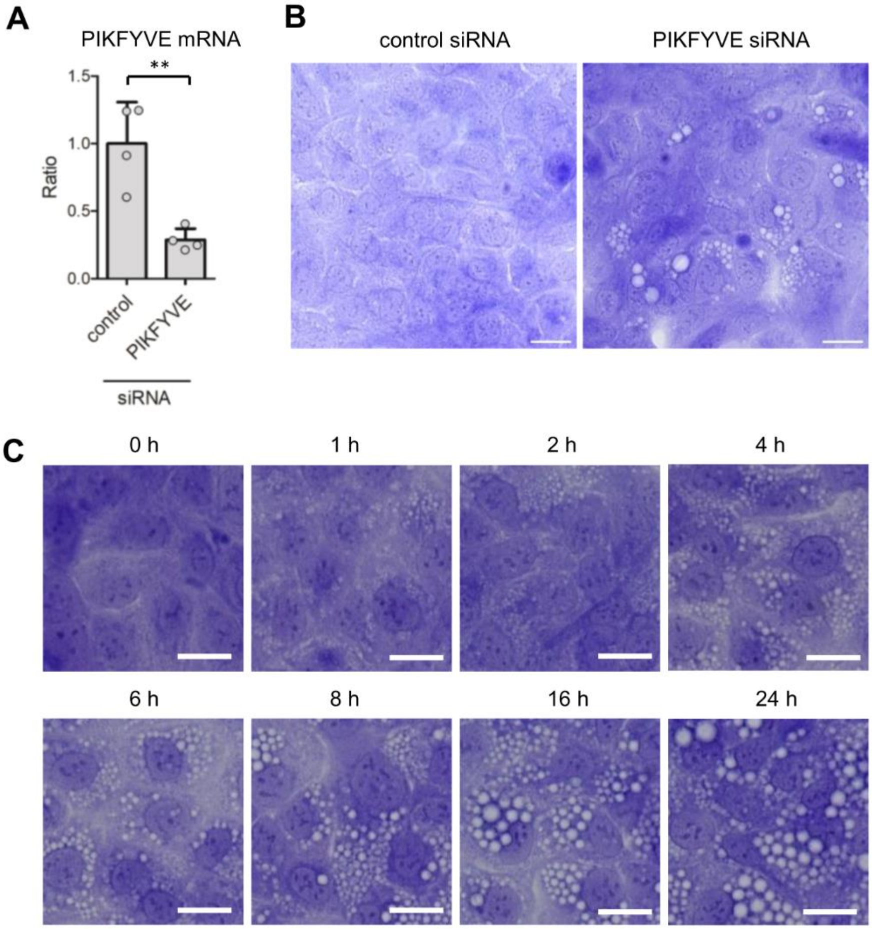


**Fig. S1. Effect of PIKFYVE inhibition on vacuole formation.** (A) Knockdown of PIKFYVE in DU145 cells was evaluated by quantitative PCR. PIKFYVE expression was normalized by the corresponding GAPDH expression. **p < 0.01 (unpaired two-tail Student's t-test; n = 4). (B) Vacuole enlargement via PIKFYVE knockdown. Representative images of crystal violet-stained cells. Bar = 20 μm. (C) Time-course of vacuole enlargement after YM201636 treatment. DU145 cells were treated with 1 μM YM201636 for the indicated times. Representative images of crystal violet-stained cells. Bar = 20 μm.

Fig. S2

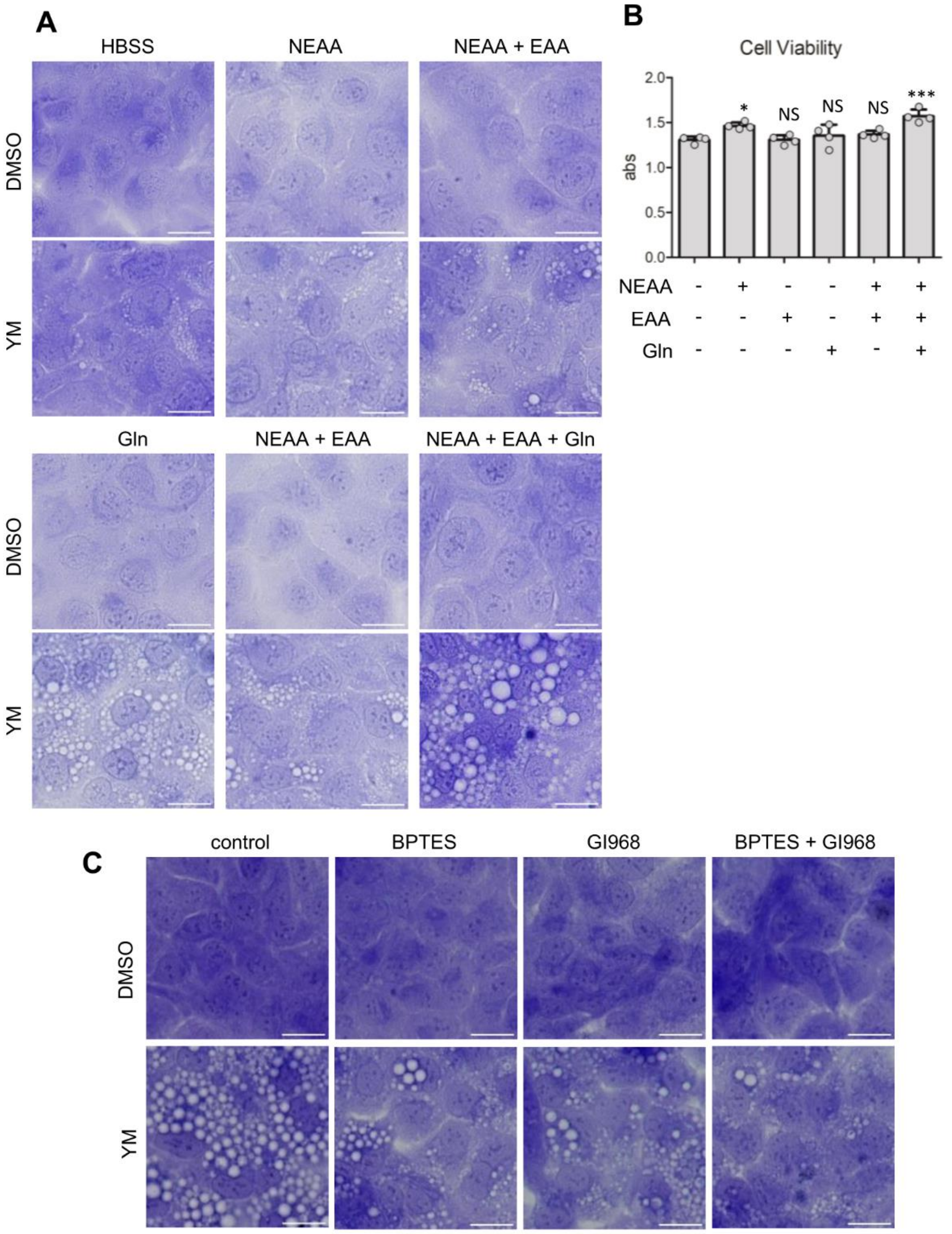


**Fig. S2. Influence of amino acids on vacuole enlargement via PIKFYVE inhibition.** (A) DU145 cells were treated with 1 μM YM for 24 h in HBSS medium with NEAAs, EAAs or glutamine (Gln). Representative images of crystal violet-stained cells. Bar = 20 μm. (B) DU145 cells were plated in a 96-well dish (20,000 cells per well) and cultured 2 days in RPMI medium supplemented with 10% FBS. Then, the medium was replaced with HBSS medium with NEAAs, EAAs or Gln and cultured for 24 h. Cell viability was evaluated by cell counting kit-8 (Dojindo, CK04) according to the manufacturer's protocol. NS, not significantly different. *$p < 0.05$ and ***$p < 0.001$ (compared to amino acid-free; n = 4). (C) DU145 cells were co-treated with 1 μM YM and 10 μM BPTES or 20 μM GI968 for 24 h in RPMI. Representative images of crystal violet-stained cells. Bar = 20 μm.

### Fig. S3

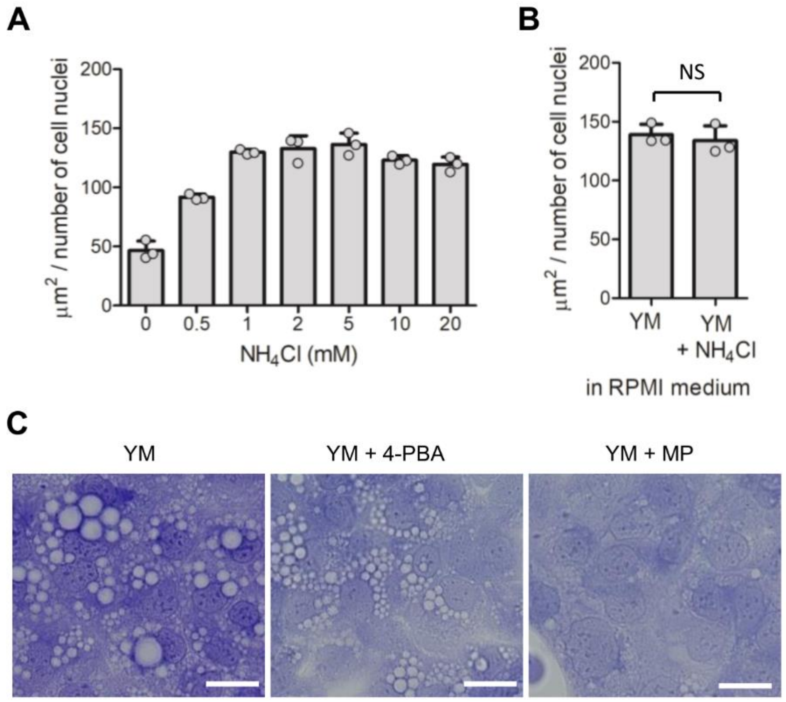


**Fig. S3. Influence of ammonia/ammonium on vacuole enlargement via PIKFYVE inhibition.** (A) DU145 cells were cultured in HBSS medium with the indicated concentration of $NH_4Cl$ and 1 μM YM for 24 h. After crystal violet staining, the areas of the vacuoles were quantified (n = 3). (B) DU145 cells were co-treated with 1 μM YM and 20 mM $NH_4Cl$ for 24 h in RPMI. After crystal violet staining, the areas of the vacuoles were quantified (n = 3). NS, not significantly different via unpaired two-tail Student's t-test. (C) Effect of ammonia scavengers on PIKFYVE inhibition-induced vacuole enlargement. DU145 cells were co-treated with 1 μM YM and 5 mM 4-phenylbutyric acid (4-PBA) or 20 mM methyl pyruvate (MP) for 24 h in RPMI. Representative images of crystal violet-stained cells. Bar = 20 μm.

### Fig. S4

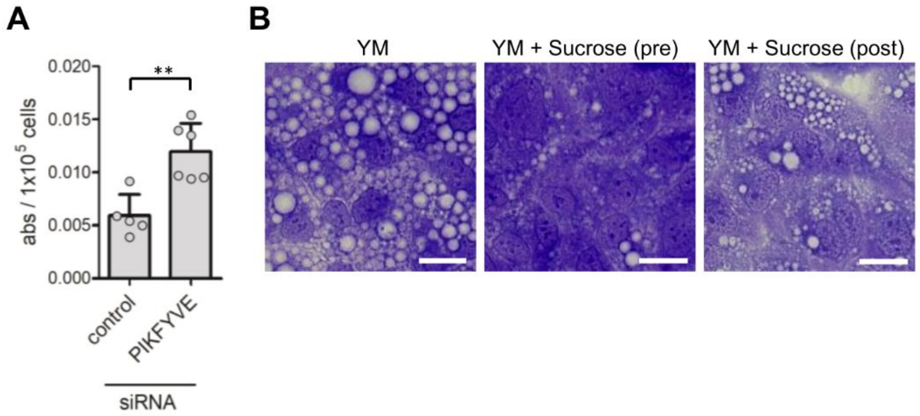


**Fig. S4. Effect of PIKFYVE inhibition on ammonium ion accumulation.** (A) DU145 cells were treated with PIKFYVE or negative control siRNA. Each cellular $NH_3/NH_4^+$ was evaluated using the Berthelot reaction. **$p < 0.01$ (unpaired two-tail Student's t-test; n = 5-6). (B) Effect of osmotic pressure on PIKFYVE inhibition-induced vacuole enlargement. DU145 cells were challenged with osmotic stress by the addition of 200 mM sucrose simultaneously with 1 μM YM (pre) or after 24 h YM treatment for 6 h (post). Representative images of crystal violet-stained cells. Bar = 20 μm.

### Fig. S5

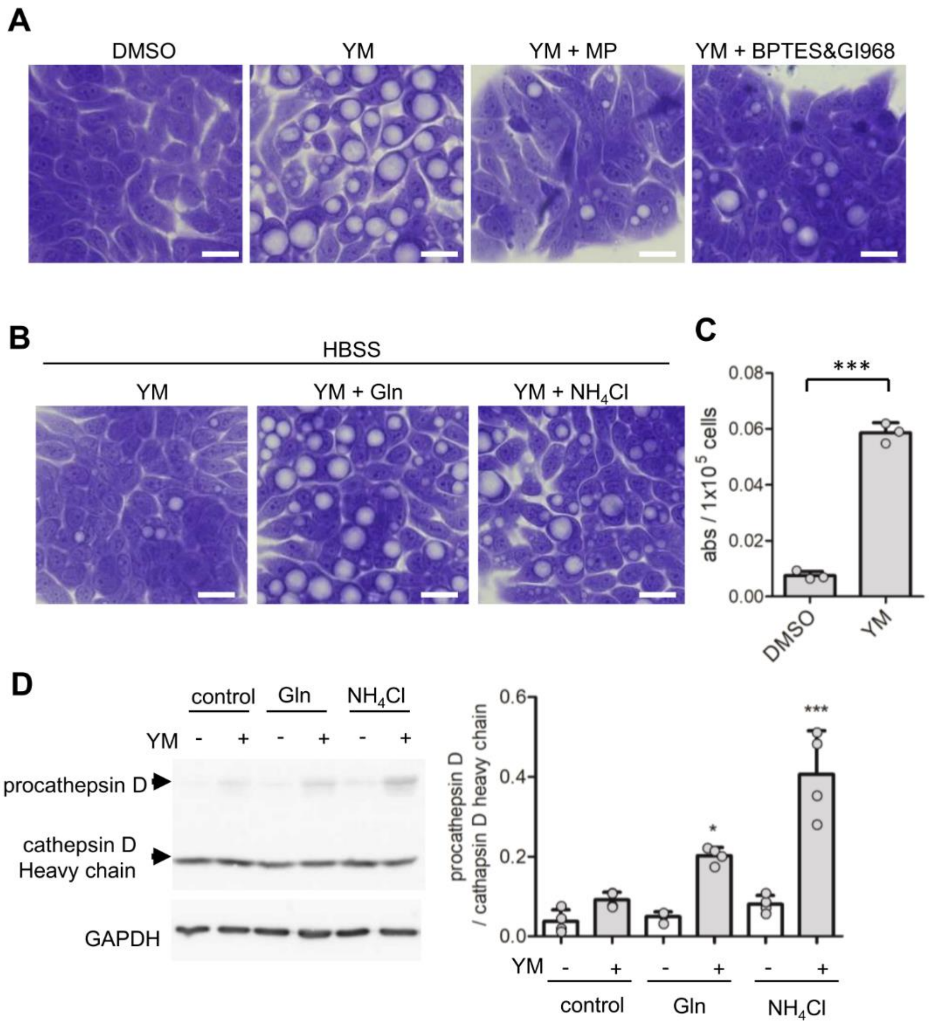


**Fig. S5. Effect of PIKFYVE inhibition on HT29 cells.** (A) HT29 cells were co-treated with 0.3 M YM201636 and 20 μM methyl pyruvate (MP) or mixture of 10 μM BPTES and 20 μM GI968 for 24 h. Representative images are shown. Bar = 20 μm. (B) HT29 cells were treated with 0.3 μM YM201636 for 24 h in HBSS medium containing 2 mM glutamine (Gln) or 2 mM $NH_4Cl$. Representative images are shown. (C) HT29 cells were treated with 0.3 μM YM201636 or DMSO (final: 0.5%) for 24 h. Each cellular $NH_3/NH_4^+$ was evaluated using the Berthelot reaction.

Values are represented as the mean and standard deviation (SD; n = 3). ***p < 0.001 (unpaired two-tail Student's *t*-test). (D) HT29 cells were treated with 0.3 μM YM for 24 h in HBSS medium containing 2 mM Gln or 2 mM $NH_4Cl$. Cell lysates were analyzed via immunoblotting using cathepsin D and GAPDH antibodies. As indicated by the arrows, procathepsin D was observed at high molecular weights, whereas the heavy chain of mature cathepsin D was observed at low molecular weights. The reduced level of mature cathepsin D was quantified as the ratio of procathepsin D to cathepsin D heavy chain (right side). *p < 0.05 and ***p < 0.001 (n = 4).

### Fig. S6

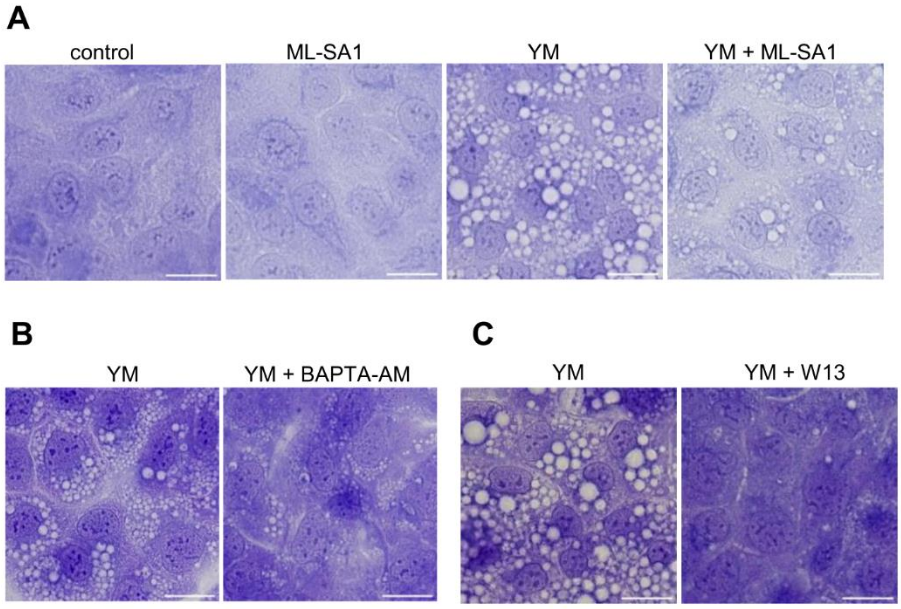


**Fig. S6. Effect of TRPML1 activator, calcium chelator and calmodulin inhibitor on PIKFYVE inhibition-induced vacuole enlargement.** (A) DU145 cells were co-treated with 1 μM YM and 20 μM ML-SA1 or DMSO (final: 0.5%) for 24 h. Representative images of crystal violet-stained cells. Bar = 20 μm. (B) DU145 cells were co-treated with 1 μM YM and 20 μM BAPTA-AM or DMSO (final 0.5%) for 6 h. (C) DU145 cells were co-treated with 1 μM YM and 20 μM W-13 or DMSO (final: 0.5%) for 24 h. Representative images of crystal violet-stained cells. Bar = 20 μm.

### Fig. S7

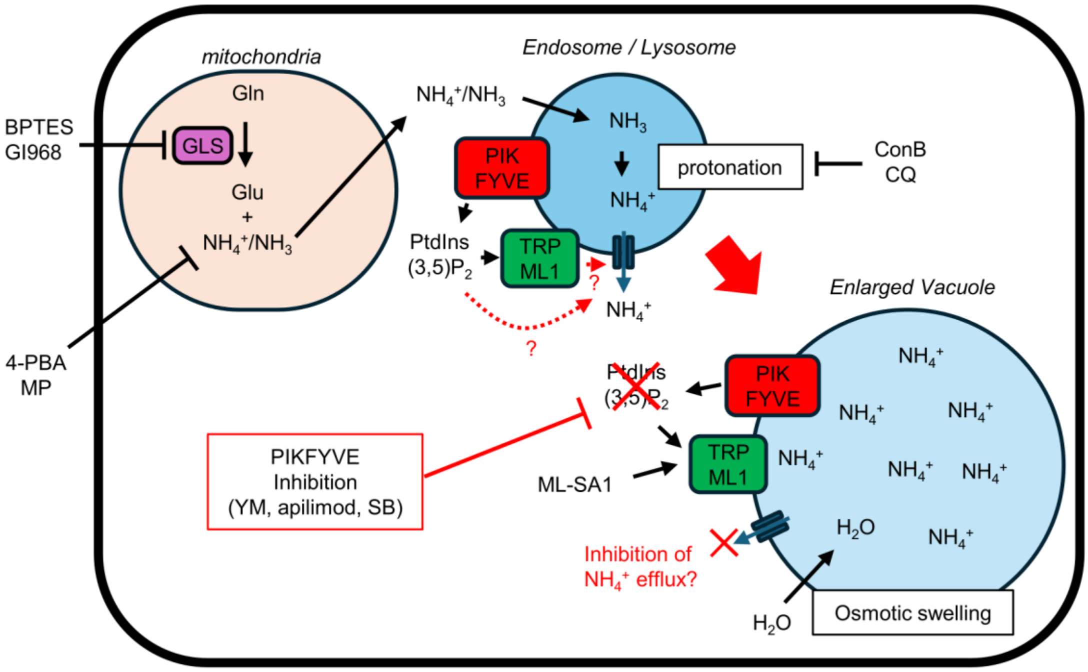


**Fig. S7. Schematic of endosomal and lysosomal enlargement upon PIKFYVE inhibition and the target of the drugs used in the study.** Glutamine (Gln) is catalyzed by glutaminases (GLS) into glutamine (Glu) and ammonium/ammonia ($NH_4^+/NH_3$) in mitochondria. $NH_3$ can enter endosomes and lysosomes and be protonated to $NH_4^+$ in the acidic environment. Efflux of $NH_4^+$ from endosomes and lysosome needs specific transporter such as SLC12A9. PIKFYVE is responsible kinase to produce PtdIns(3,5)$P_2$ on endosomes and lysosomes. PtdIns(3,5)$P_2$ affects membrane protein functions, such as TRPML1 activation. PIKFYVE inhibition leads to enlargement of endosomes and lysosomes via $NH_4^+$ accumulation-induced osmotic swelling. This suggests that PIKFYVE activity is required for $NH_4^+$ efflux. We hypothesize that PtdIns(3,5)$P_2$ regulate $NH_4^+$ efflux transporter activity directly or indirectly via TRPML1 and others (red dashed arrow). In this study, PIKFYVE was inhibited by YM201636

(YM), apilimod and SB202190 (SB). BPTES and glutaminase inhibitor compound 968 (GI968) inhibit Gln metabolism by GLS, while Sodium 4-phenylbutyrate (4-PBA) and methyl pyruvate (MP) scavenge ammonia. Thus, these drugs inhibit vacuole enlargement by the PIKFYVE inhibitor. Concanamycin B (ConB) and chloroquine (CQ) reduce protonation of $NH_3$ through lysosomal neutralization as an inhibitor of proton pump V-ATPase and as a weak base, respectively. TRPML1 can be activated by mucolipin synthetic agonist 1 (ML-SA1) instead of PtdIns(3,5)$P_2$. ML-SA1 suppresses PIKFYVE inhibition-induced $NH_4^+$ accumulation and vacuole enlargement.

### Fig. S8

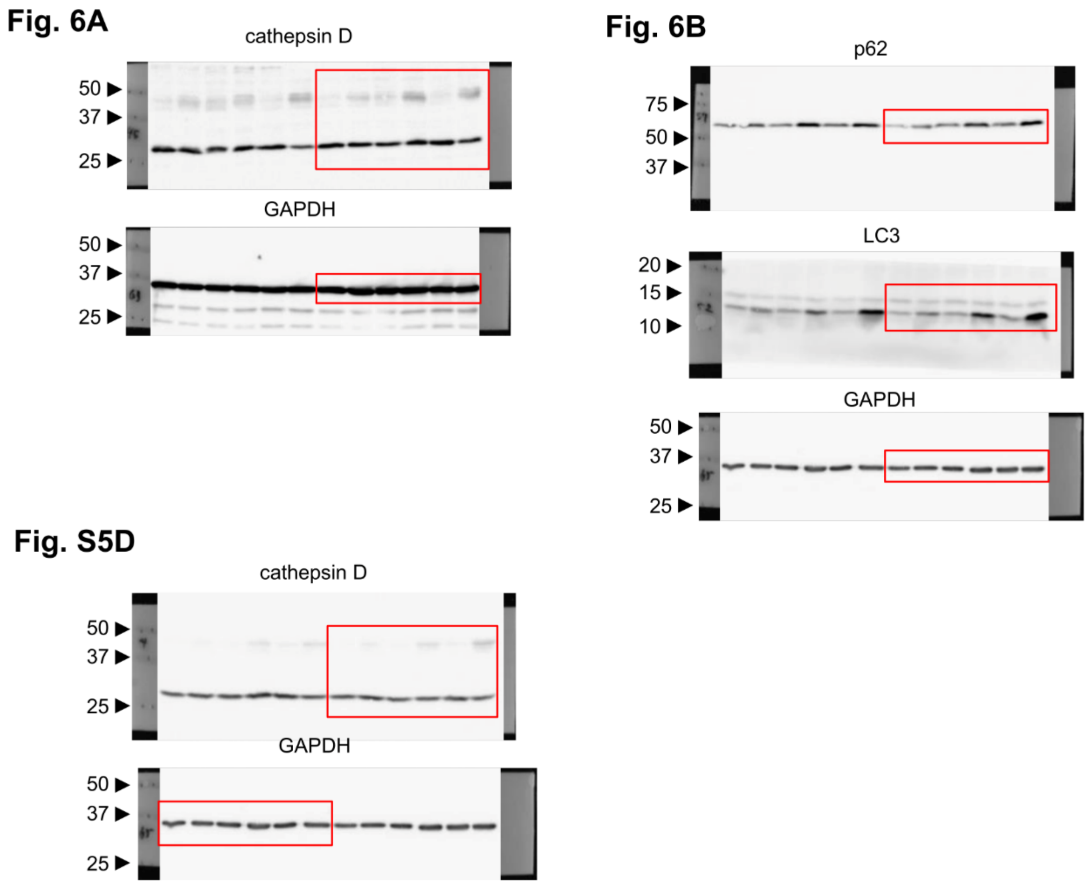


**Fig. S8. Western blot transparency.** The edge of each membrane is shown by displaying the composite brightfield image to the left and right. The red box indicates the area trimmed for the figure. GAPDH results in Fig. 6B and Fig. S5D were taken from the same Western blot result.
